# AI inspired discovery of new biomarkers for clinical prognosis of liver cancer

**DOI:** 10.1101/2022.05.03.490448

**Authors:** Junhao Liang, Weisheng Zhang, Jianghui Yang, Meilong Wu, Qionghai Dai, Hongfang Yin, Ying Xiao, Lingjie Kong

## Abstract

Tissue biomarkers are crucial for cancer diagnosis, prognosis assessment, and treatment planning. However, few of current biomarkers used in clinics are robust enough to show a true analytical and clinical value. Thus the search for additional tissue biomarkers, including the strategies to identify them, is imperative. Recently, the capabilities of deep learning (DL)-based computational pathology in cancer diagnosis and prognosis have been explored, but the limited interpretability and generalizability make the results difficult to be accepted in clinical practice. Here we present an interpretable human-centric DL-guided framework—PathFinder (Pathological-biomarker-finder)— that can inspire pathologists to discover new tissue biomarkers from well-performing DL models, which bridges the gap between DL and clinical prognosis. By combining sparse multi-class tissue spatial distribution information of whole slide images (WSIs) with attribution methods, PathFinder can achieve localization, characterization, and verification of potential biomarkers, while guaranteeing state-of-the-art prognostic performance. With the inspiration of PathFinder, we discovered that tumor necrosis in liver cancer, a long-neglected factor, has a strong relationship with patient prognosis. Thus we proposed two clinically independent indicators, including necrosis area fraction and tumor necrosis distribution, for practical prognosis, and verified their potentials in clinical prognosis according to Reporting Recommendations for Tumor Marker Prognostic Studies (REMARK)-derived criteria. Our work demonstrates a successful example of introducing artificial intelligence (AI) into clinical practice in a knowledge discovery way, which can be adopted in identifying biomarkers in various cancer types and modalities.

## Introduction

Pathological analysis of WSIs is the gold standard for cancer diagnosis and prognosis. Tumor classification, staging, and prognosis are assessed according to tissue biomarkers on WSIs^1,2^. Unfortunately, even though various tissue biomarkers have been proposed, few of them is robust with high sensitivity and specificity^3,4^. Thus there is still a desperate need for identifying additional robust biomarkers to guide tumor diagnosis and prognosis, and to direct the research of tumor mechanism^5–7^. Specifically in cancer prognosis, with the advancement of computational pathology in recent years, DL models based on end-to-end training can predict a risk score that outperforms current clinical staging, showing the potential of learning knowledge from current medical data^8–12^. However, due to limited interpretability and generalizability, DL-based risk score is still difficult to be accepted as a useful biomarker for clinical prognosis^6,13,14^.

Considering that clinicians are likely to keep playing the central role in patient care, it is essential to focus the development and evaluation of AI-based clinical algorithms on their potential to augment rather than replace human intelligence^15–17^. Although some studies have attempted to use established biomarkers and attribution methods to verify the credibility of abstract risk scores^8–11^, this strategy fails in generating new knowledge for clinical prognosis. Knowledge discovery based on AI, especially the discovery of new or dominant prognostic biomarkers of clear pathological significance and explicit mathematical model, will open up new direction of human-centric AI for cancer prognosis.

Different from that in the fields of genetics where biologically informed sparse DL models combined with attribution methods has been used to guide preclinical discovery^18^, the identifying of tissue biomarkers from well-performing prognostic DL models is challenging^8–12^. On one hand, the input multi-dimensional images of WSIs for prognosis are of high information density, compared to genetics inputs which are usually one-dimensional vector and have specific labels or descriptions. Thus it is difficult to build a sparse network while guaranteeing the prognostic performance^19^. On the other hand, current attribution methods usually achieve a two-dimensional attribution map for spatial attribution positioning^13,14^, which is far from locating specific high-attribution features in high-information-density input. These two problems lead to insufficient interpretation, as low-dimensional attribution knowledge is used to interpret abstract results based on high-dimensional inputs. Even worse, it makes one use pre-existing knowledge in explanation, which contradicts the aim of discovering new biomarkers^19–21^.

Histologically, gigapixel WSIs can be regarded as self-multimodal information sources with both slide-level macro mode and region-level micro mode^14^. The former contains multi-class tissue spatial distribution and interaction information, while the latter contains cell texture and structure information. However, limited by GPUs (Graphics Processing Units) memory and deep neural network architecture, WSIs are generally cut into patches and only the micro mode information is paid attention to in most DL-based studies^9,10,22–24^. Moreover, in clinics, due to the lack of precise quantification of WSIs, the relationship between tissue spatial distribution and patients’ prognostic result is still not clear.

Here we propose an interpretable, human-centric, DL framework, named as PathFinder, that uses the sparse multi-class tissue spatial distribution information of WSIs for assessing prognosis and discovering new biomarkers. Using the macro mode of WSIs, which is of low information density that perfectly matches current spatial-positioning attribution methods, Pathfinder can achieve state-of-the-art prognostic performance. Inspired by the exact and intuitive attribution maps of PathFinder, we found tumor necrosis in liver, a common but overlooked tissue, has a strong relationship with patients’ prognosis, based on which we characterized two significant indicators for clinical prognosis.

## Results

### Interpretable AI-based framework for macro biomarker discovery

Figure 1 shows the workflow of Pathfinder. It consists of three parts: macro mode acquiring, prognostic deep neural network training, and new biomarker discovery. We first trained the multi-class tissue segmentation network PaSegNet to obtain the multi-class tissue probability heatmaps as the macro mode of WSIs (Methods). In order to acquire high-quality macro mode, we proposed meta annotation, a data-centric annotation method that combined with pathological priors to bridge the gap between current pathological annotation methods and DL training requirements, and achieved efficient, high diversity, and low similarity class-balanced training dataset (Methods, Extended Data Fig. 1). With the macro mode of WSIs, we built MacroNet for high-precision prognosis, which is composed of a convolution feature extractor and a multilayer perceptron (MLP) with a batch normalize layer^25^ (Methods). Using only time-to-event patient death information as the input mode label and Cox proportional likelihood loss as the network loss, the MacroNet can learn to predict the patients’ risk score based on macro mode only. Then we used attribution methods on the trained MacroNet to acquire the attribution map of input image^26^, and overlapped the attribution map on the corresponding multi-class segmentation map. The generated two-dimensional attribution map shows the spatial areas that MacroNet focuses on, which matches well with the sparse multi-class tissue spatial distribution information, making the interpretation more direct and objective. Based on integrative analysis of macro mode and attribution map, pathologists can propose the hypothesis of the biomarkers that the model is concerned with, followed by quantitatively characterization. The new biomarkers, whose visualizations are similar with the corresponding attribution map, were used as indicators to perform multivariate analysis according to REMARK-derived criteria^27^. After testing with clinical dataset, new biomarkers of significantly independent prognostic effect were discovered.

**Fig. 1.**
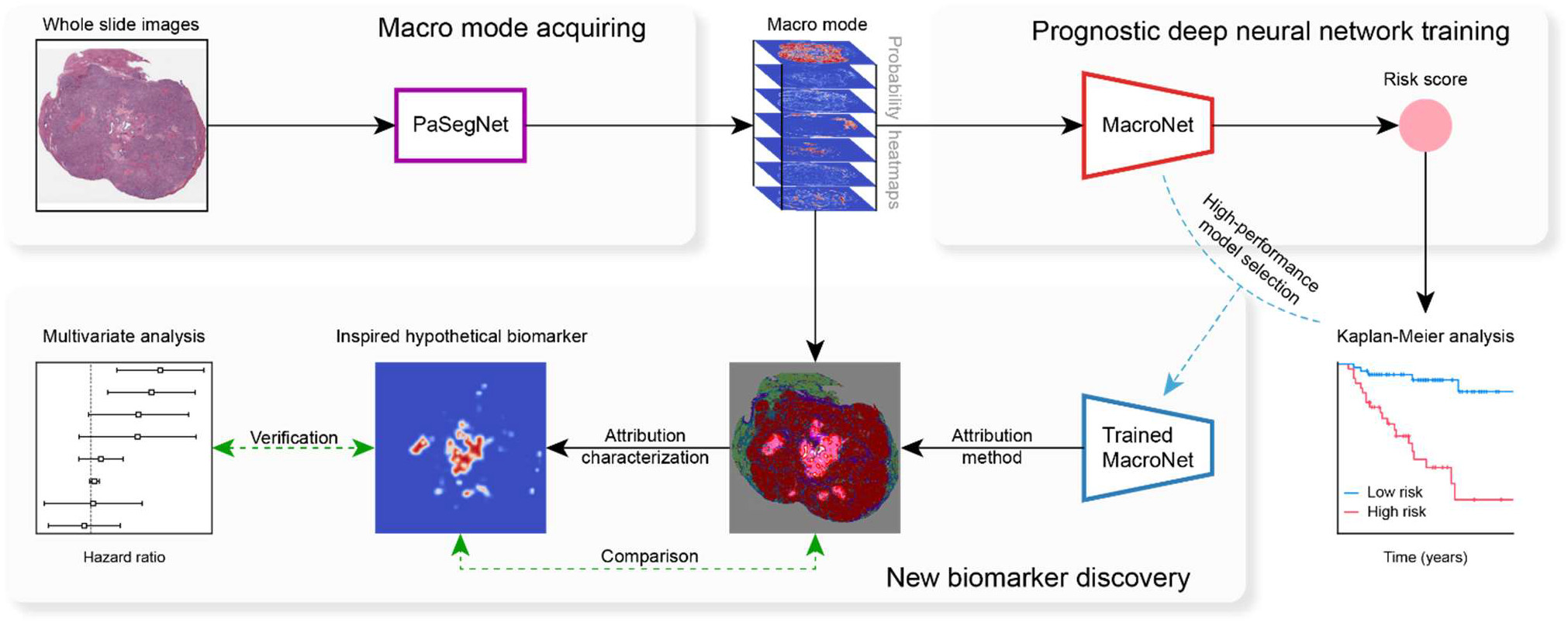
The workflow of PathFinder. Digitized high-resolution histology slides of patients serve as the input into the framework. The WSI is first processed with PaSegNet, a convolutional neural network, to obtain the spatial distribution probability heatmaps of 7 common liver tissues. The achieved macro mode and the corresponding survival time are used as the image-label pair to train the MacroNet, a prognostic convolutional neural network with the output of corresponding risk score for guiding the patient’s prognosis. Then one can apply the attribution method to the trained, well-performing MacroNet to explore the model’s spatial focus area, from which to get the inspiration of potential prognostic biomarkers. Following that, these hypothetical biomarkers are modeled based on the macro mode to achieve quantification and characterization, in which the ones similar to the attribution map after visualization are selected as candidate biomarkers and used as indicators for multivariate analysis. After testing with clinical dataset, the significantly independent prognostic indicators can be identified.

With Pathfinder, we performed the discovery of new tissue biomarkers for clinical prognosis of hepatocellular carcinoma (HCC), which is the fourth leading cause of cancer-related death worldwide^28^. In this study, we collected 342 WSIs from 330 patient samples in The Cancer Genome Atlas Liver Hepatocellular Carcinoma dataset (TCGA dataset) and 1182 WSIs from 83 patient samples in Beijing Tsinghua Changgung Hospital dataset (QHCG dataset) (Extended Data Figs. 2, 3). As for the case that there are multiple WSIs for a patient, we selected the one of largest tumor fraction as the patient’s representative WSI, as discussed later. We trained MacroNet in a 10-fold cross-validation on TCGA dataset, tested the generalization of trained model on QHCG dataset. In order to better compare the prognostic performance of MacroNet, we also designed and trained MicroNet and M2MNet for prognosis task. The former one is based on micro mode, and the latter one is based on both macro mode and micro mode (Methods).

### Evaluation of model performance

We first evaluated the multi-class classification performance of PaSegNet on the internal test set of QHCG dataset and external independent test sets including TCGA dataset and Pathology AI Platform 2019 challenge dataset (PAIP dataset). Confusion matrices and receiver operating characteristic (ROC) curves are used to demonstrate classification results (Fig. 2a, Extended Data Fig. 4). The macro-average accuracy and area under the curve (AUC) are selected to evaluate model performance. Across all test sets, PaSegNet achieved accuracy of 0.948, 0.956, 0.941, and AUC of 0.9980, 0.9984, 0.9974, on QHCG, TCGA, PAIP test set, respectively. The results show that the PaSegNet trained on the meta-annotated dataset can achieve accurate multi-class tissue classification. To evaluate the segmentation performance of WSIs, we further visualized the multi-class tissue probability heatmaps and segmentation maps obtained by PaSegNet, both of which demonstrate that the model can accurately and smoothly segment WSIs and identify small key lesion areas (Extended Data Fig. 5). In general, PaSegNet trained on the meta-annotation dataset can efficiently quantify WSIs’ macro mode and ensure the following prognostic network training.

**Fig. 2.**
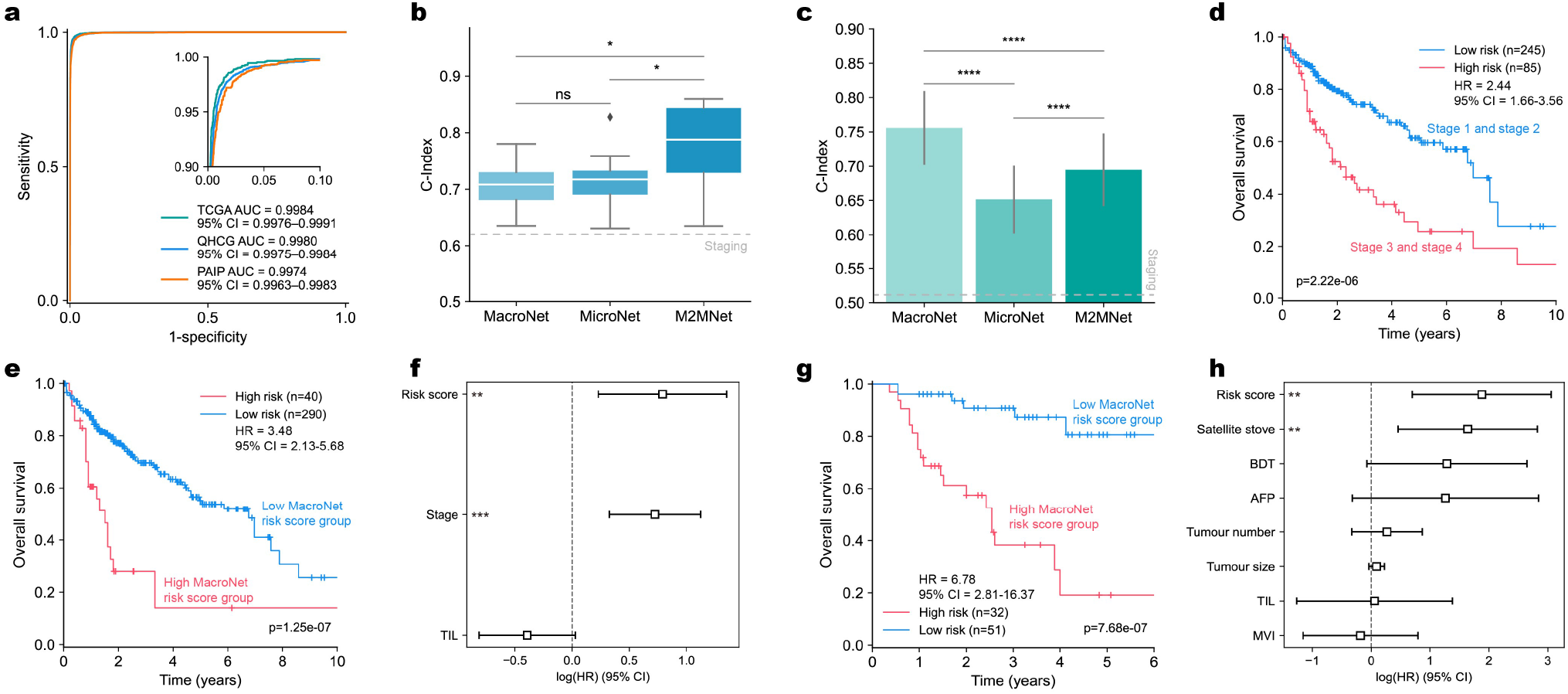
Performance of Pathfinder in the discovery of new tissue biomarkers for clinical prognosis of HCC. **a**, ROC curves for the multi-class tissue classification, evaluated on the internal test set (QHCG) and external independent test sets (TCGA, PAIP). CI, confidence interval. **b**, C-Index distribution of MacroNet, MicroNet, and M2MNet on TCGA dataset in a 10-fold cross-validation. Boxes indicate quartile values and whiskers extend to data points within 1.5× the interquartile range. **c**, C-Index performance of MacroNet, MicroNet, and M2MNet on QHCG test set. Error bars indicate 95% confidence intervals. **d**, Kaplan-Meier analysis of patient stratification of clinical staging patients. **e, g**, Kaplan-Meier analysis of patient stratification of low and high-risk patients via MacroNet on TCGA dataset (**e**) and QHCG dataset (**g**), respectively. Logrank test is used to test statistical significance in survival distributions between low-risk and high-risk patients. **f, h**, Multivariable analysis of factors associated with overall survival and MacroNet risk score on TCGA dataset (**f**) and QHCG dataset (**h**), respectively. HR, hazard ratio; Stage, AJCC staging; TIL, tumor infiltrating lymphocytes digital score; BDT, bile duct thrombosis; AFP, alpha-fetoprotein; MVI, microvascular invasion. The results of univariate and multivariate analyses are described in details in Extended Data Tables 1, 2. The significance level shown in **b, c** is determined using a two-sided Mann-Whitney-Wilcoxon test (**b**) and a two-sided two-sample *t*-test (**c**). \**p* < 0.05, \*\**p* < 0.01, \*\*\**p* < 0.001, \*\*\*\**p* < 0.0001.

We next evaluated the prognostic capability of MacroNet, MicroNet, and M2MNet, by using 10-fold cross-validation on TCGA dataset. To compare the performance of prognostic networks, we used the median of cross-validated concordance index (C-Index) to measure the predictive accuracy of each model, Kaplan-Meier curves to visualize the quality of patient stratification between predicted high-risk and low-risk patients, and the logrank test to test the statistical difference between high-risk and low-risk groups. MacroNet achieved a C-Index of 0.708, similar to the C-Index 0.717 using MicroNet and lower than the C-Index 0.787 using M2MNet (Fig. 2b). In visualizing the Kaplan-Meier survival curves of predicted high-risk and low-risk patient groups, MacroNet also showed well discrimination between the two risk groups (p-value = 1.25×10^−7^) compared to M2MNet and clinical staging (Figs. 2d, e, Extended Data Fig. 7a).

We further evaluated the models’ generalization capability by training the models on TCGA dataset and testing them on QHCG dataset. MacroNet achieved a C-Index of 0.754, whereas M2MNet and MicroNet achieved C-Indices of 0.695 and 0.652, respectively (Fig. 2c). In addition, the Kaplan-Meier survival curves of MacroNet showed well discrimination between two risk groups (p-value = 7.68×10^−7^) on QHCG dataset, as M2MNet did (Fig. 2g, Extended Data Fig. 7b). Furthermore, the multivariable analysis revealed that the risk score predicted by MacroNet (Hazard ratio (HR): 2.21, 95% confidence interval (CI): 1.26 to 3.86, p-value = 0.0057, TCGA dataset; HR: 6.56, 95% CI: 2.01 to 21.36, p-value = 0.0018, QHCG dataset) was independent of other clinicopathological characteristics (Figs. 2f, h, Extended Data Tables 1, 2). These results indicate that MacroNet can achieve state-of-the-art prognostic performance using only macro mode of WSIs and has potential in finding useful prognostic biomarkers.

### Discovery, characterization, and verification of new tissue biomarkers

In order to interpret why MacroNet can achieve high-performance prognosis and to explore which macro features largely contribute to risk score, we conducted an integrated analysis from both global and individual perspectives. We counted the difference in the tissue fractions in patients of high-risk scores and low-risk scores from a global perspective, and found that the necrosis fraction is significantly higher in the high-risk score group (Extended Data Figs. 6a, c). Then we analyzed the segmentation map of high-risk and low-risk WSIs, and observed that necrosis occurred in every high-risk WSI, but not in all low-risk WSIs (Fig. 3a). From an individual perspective, we used the attribute method to locate the areas where MacroNet focused on in the form of a two-dimensional heatmap, and overlapped the result with the segmentation map for better visualization (Fig. 1). We discovered that the areas of high contribution are almost the junctions of necrosis and other tissues (Fig. 3b), which is consistent with our former conclusions obtained from the global perspective. All the discoveries inspired us that necrosis may have a strong relationship with HCC prognosis.

**Fig. 3.**
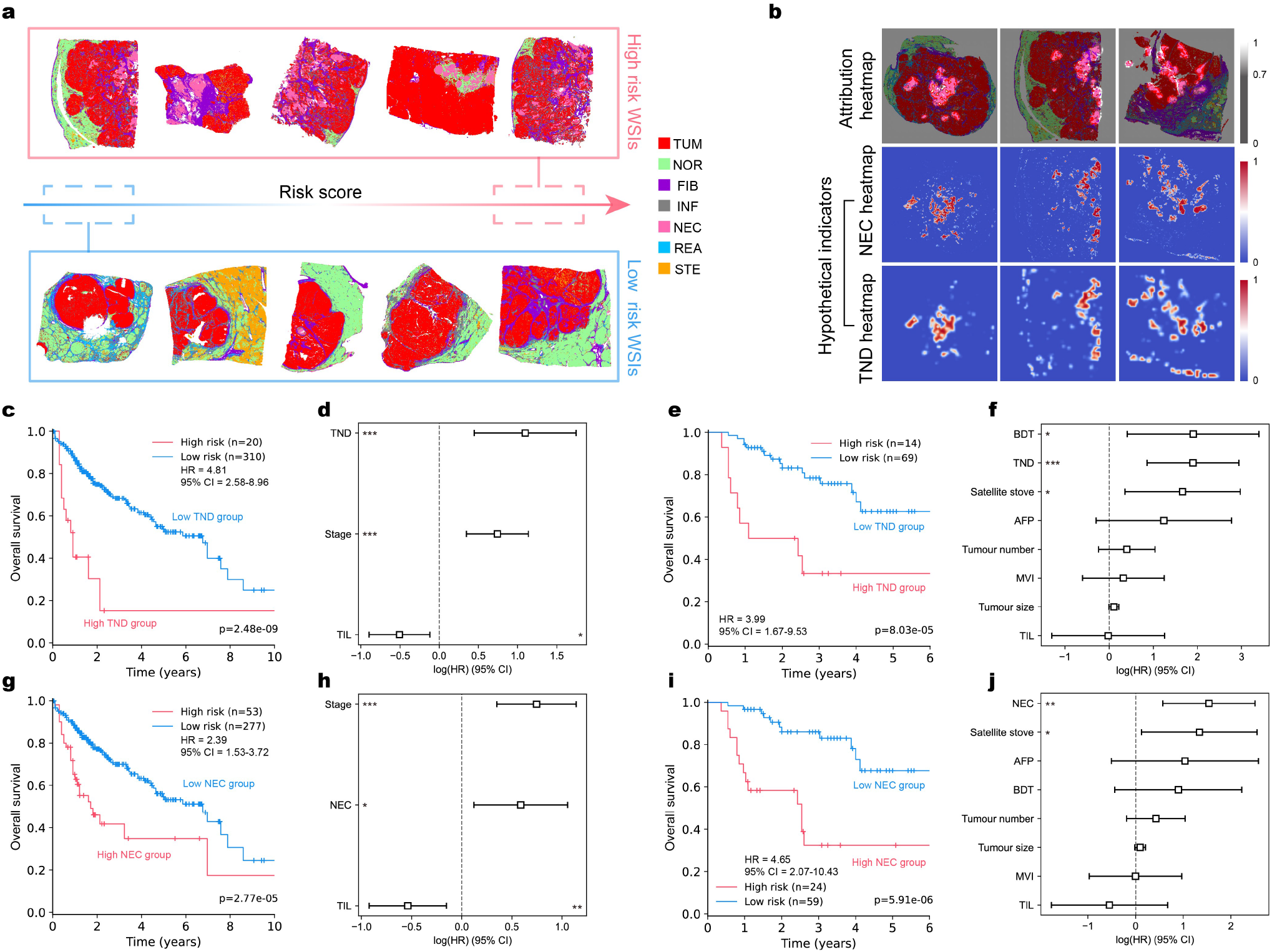
Discovery, characterization, and verification of new tissue biomarkers. **a**, Segmentation maps of low and high-risk WSIs predicted by MacroNet on TCGA dataset and QHCG dataset. **b**, Attribution heatmaps of WSIs segmentation maps and their corresponding visualization results of NEC and TND hypothetical indicators. **c, e**, Kaplan-Meier analysis of patient stratification of low (low TND score) and high-risk (high TND score) patients on TCGA dataset (**c**) and QHCG dataset (**e**). **d, f**, Multivariable analyses of TND and other factors associated with overall survival on TCGA dataset (**d**) and QHCG dataset (**f**). **g, i**, Kaplan-Meier analysis of patient stratification of low (low NEC score) and high-risk (high NEC score) patients on TCGA dataset (**g**) and QHCG dataset (**i**). **h, j**, Multivariable analyses of NEC and other factors associated with overall survival on TCGA dataset (**h**) and QHCG dataset (**j**). **c, e, g, i**, The results of logrank tests, showing statistical significances in survival distributions between low and high risk patients. **d, f, h, j**, The results of univariate and multivariate analyses. Details are shown in Extended Data Tables 1, 2. HR, hazard ratio; Stage, AJCC staging; TIL, tumor infiltrating lymphocytes digital score; BDT, bile duct thrombosis; AFP, alpha-fetoprotein; MVI, microvascular invasion; TUM, tumor; Nor, normal; FIB, fibrosis; INF, inflammation; NEC, necrosis; REA, bile duct reaction; STE, steatosis. \**p* < 0.05, \*\**p* < 0.01, \*\*\**p* < 0.001.

To make the DL-based MacroNet acceptable in clinical practice, we proposed two hypotheses of new biomarkers, namely necrosis area fraction in WSIs (NEC) and tumor necrosis distribution (TND), based on above integrated analyses and inspirations of MacroNet. We first established mathematical models of these two indicators to characterize them, and achieved their quantification based on the existing macro mode (Methods). By visualizing these two indicators and comparing them with the corresponding attribution map, we found that these two hypothetical indicators can well characterize the features that MacroNet pays attention to, indicating that these two clinically available indicators are of great potential to affect the prognosis of the risk score given by MacroNet (Fig. 3b). It also should be noted that these biomarkers are objective and universal pathological features, considering that NEC is a common and inherent attribute of WSIs, and TND is a newly designed indicator that takes into account the spatial distribution and interaction between tumor and necrosis.

To verify whether NEC and TND are independent prognostic indicators, we investigated the prognostic significance of these two indicators on both TCGA and QHCG datasets using Kaplan-Meier curves and Cox hazard analysis by conducting univariate and multivariate analyses of clinicopathological parameters. Additionally, to compare the performance with new clinical indicators inspired by AI, we quantified tumor-infiltrating lymphocytes (TILs), which is already known as a prognostic factor and is significantly different between high-risk group and low-risk group (Extended Data Figs. 6b, c, Methods)^12,29^, as an indicator designed based on known clinical experience. The Kaplan-Meier curves and logrank test based p-values showed that NEC and TND can significantly distinguish high-risk and low-risk groups on both TCGA and QHCG datasets (Figs. 3c, e, g, i). The univariate and multivariable analyses revealed that the dependences of overall survival on NEC (HR: 4.66, 95% CI: 1.77 to 12.28, p-value = 0.0019, QHCG dataset; HR: 1.80, 95% CI: 1.13 to 2.87, p-value = 0.0133, TCGA dataset) and TND (HR: 6.67, 95% CI: 2.36 to 18.85, p-value = 0.0003, QHCG dataset; HR: 3.00, 95% CI: 1.56 to 5.74, p-value = 0.0009, TCGA dataset) were more significant than most clinical indicators including TILs (Figs. 3d, f, h, j). This suggests that the two indicators are independent of other clinicopathological characteristics. In addition, NEC (HR: 3.31, 95% CI: 1.73 to 6.30, p-value = 0.0003) and TND (HR: 2.92, 95% CI: 1.52 to 5.60, p-value = 0.0012) can even be used as significant indicators in recurrence prediction (Extended Data Figs. 7c-i, Extended Data Table 3). It is worth noting that the Cox’s proportional hazard model was able to achieve a C-Index 0.7 without utilizing additionally clinical variables or risk score predicted by DL methods, as it makes predictions only based on NEC (C-Index: 0.703) or TND (C-Index: 0.691) (Figs. 4d, e).

**Fig. 4.**
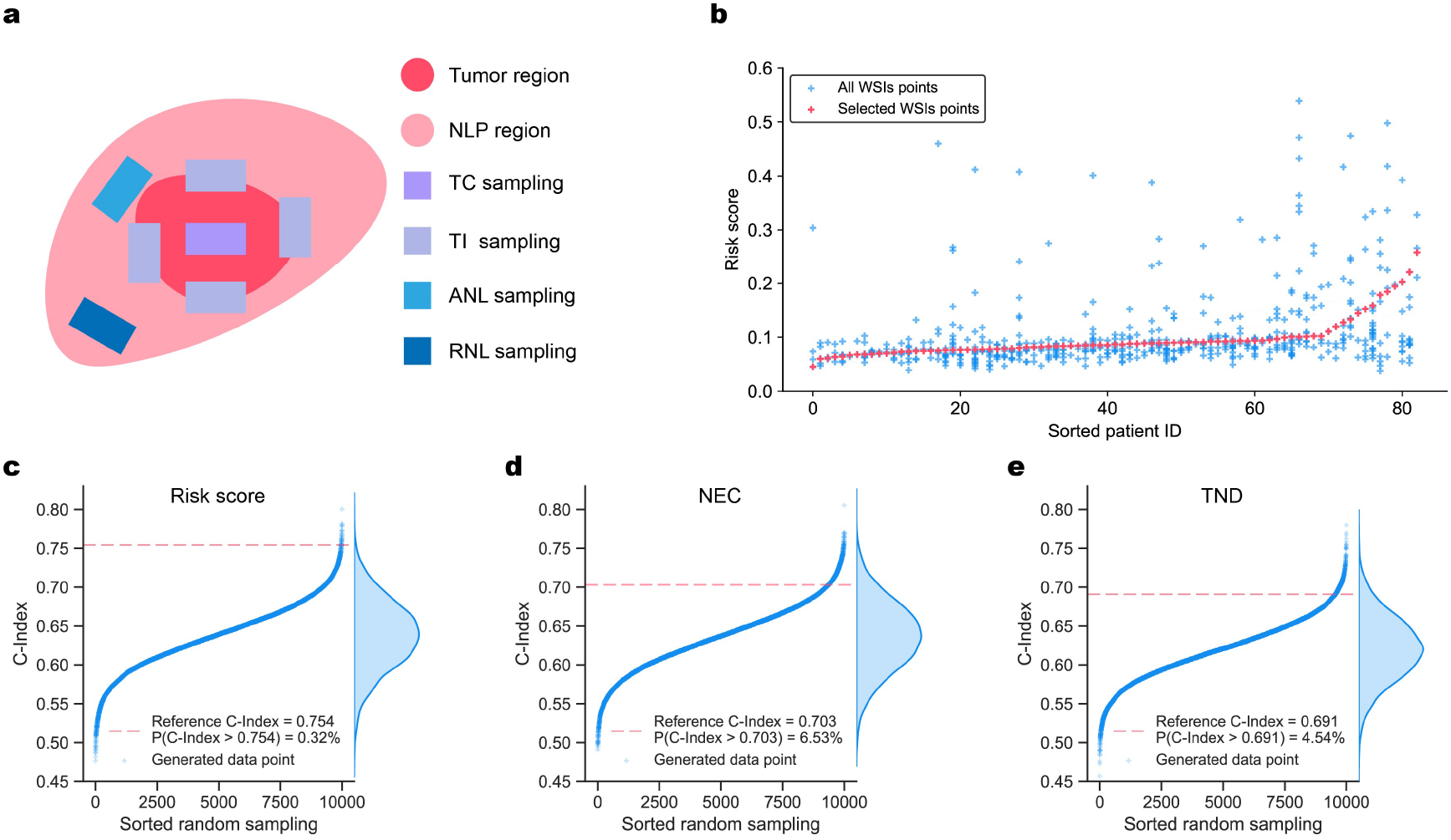
Exploring the robustness of macro mode indicators. **a**, Sampling strategy of clinical WSIs. NLP, non-neoplastic liver parenchyma; TC, tumor center; TI, tumor-liver interface; ANL, adjacent non-neoplastic liver; RNL, remote non-neoplastic liver. **b**, Deviations in the risk scores predicted by MacroNet from different WSIs of a patient. The risk scores of all WSIs (excluded WSIs without tumor) of 83 patients are ranked in ascending order based on the selected WSI points. Each patient has more than one WSIs points (blue points on a specific abscissa), in which the selected WSIs to characterize the patient’s final risk score is labelled as red points. **c-e**, Random selection strategy simulations of MacroNet risk score (**c**), NEC (**d**), and TND (**e**), respectively. The red dotted lines represent C-Indices of MacroNet risk score, NEC, and TND under the largest tumor fraction selection rule. Each blue point represents the C-Index of one random selection simulation, and all the blue points are ranked in ascending order based on their C-Indices. The distribution of these points with respect to the C-Index is shown on the right side of the image.

Overall, the above results verified necrosis as a new biomarker for prognosis. We demonstrated that the prognostic performance of the AI inspired indicators based on WSIs macro mode is comparable to the performances of various DL models based on WSIs micro mode, genomics, and multimodality^9–12^.

### Robustness of macro mode indicators

In clinical practice, there are generally many WSIs with different sampling positions from a patient (Fig. 4a). As the micro mode is not greatly affected by the sampling locations, the prognostic DL models trained on the micro mode rarely discuss the situation where a patient has multiple WSIs^8^. However, different sampling positions will cause huge differences in the macro mode, which will lead to deviations in the risk scores predicted by MacroNet (Fig. 4b). Exploring how to select representative WSI from multiple WSIs of a patient becomes an unavoidable problem in applying macro indicators in clinical prognosis.

In our former study, we selected the largest tumor fraction one as the patient’s representative WSI. In order to explore the robustness and effectiveness of this selection rule in clinical prognosis, we calculated the risk score, TND, and NEC of all WSIs, and randomly selected one from the multiple WSIs of a patient as the representative WSI, with C-Index being used to measure the accuracy of prognosis under this random sampling standard. After 10,000 simulations under random selection strategy, the prognostic performance of our former selection rule is better than most random selections (Figs. 4c, d, e). Even for NEC and TND, two objective and universal biomarkers, the results based on largest tumor fraction selection rule were better than 94% of the results based on random selection rule, indicating that the largest tumor fraction selection rule can be adopted with NEC and TND biomarkers for clinical prognosis.

## Discussion

We present PathFinder as a complete framework of AI inspired discovery of clinically acceptable biomarkers. Instead of using DL to predict a risk score from WSIs^8–11,24^, we focus on proposing human-centric workflows for inspiring pathologists to discover new clinically acceptable biomarkers from well-performing black-boxes. We show a new method of bridging AI and clinical prognosis, and prove the potential of AI in learning and exploring new prognostic biomarkers based on large datasets and objective survival information.

To overcome the limited interpretability and generalizability of DL-based risk scores, we simplified the input of DL models and explored the relationship between multi-class tissue spatial distribution and prognosis, for the first time. Different from utilizing pre-trained networks to compress WSIs^8,10,24,30^, our input is more sparse and has explicit medical meaning, which enables the attribution method to characterize the biomarkers, that the model focuses on, more accurately. Our results show that the prognostic performance of DL is still good even when the input is reduced from WSIs of several gigabytes to macro mode of several megabytes. This indicates that the multi-class tissue spatial distribution of WSIs has prognostic information and the conventional inputs of prognostic DL models are redundant.

In this study, we did not target AI as a substitute for pathologists, but as a tool for pathologists to mine dominate biomarkers. Just as AI guides mathematical intuition^31^, pathologists can formulate specific hypotheses based on their clinical experience, and then use PathFinder to deeply mine the connection between hypotheses-relevant information and prognosis. Inspired by PathFinder, we defined two necrosis-related clinical prognostic indicators, NEC and TND, and demonstrated their feasibility in HCC prognosis. Even as a common tissue in liver cancer, necrosis has caught few attentions and has not been put into clinical staging guidelines in detail^32–34^. Our findings demonstrate that AI can analyze data more objectively and alert us about missing information. Different from highly diverse tumor tissues, necrosis is easier to be distinguished in both clinics and computer vision, which makes it convenient for clinical prognosis. Meanwhile, the mechanisms between tumor and necrosis are still unclear. The significant effect of TND and NEC on prognosis may suggest that the spatial distribution of tissue is worth considering in researches of necrosis mechanisms. Additionally, tumor necrosis is postulated to be caused by tumor necrosis factors^35^, which have been found significant correlations with TILs^36,37^. However, our results suggest a low correlation between tumor necrosis and TILs (Extended Data Figs. 7j, k), indicating that HCC necrosis may have its own specific causes and mechanisms.

As products of knowledge discovery, TND and NEC have clear pathological significance and explicit mathematical model. The strong generalizability of these new biomarkers is evaluated on TCGA and QHCG datasets, suggesting the great advantages of human-centric AI for knowledge discovery and clinical prognosis.

Same as all commonly used DL models, the focusing features of PathFinder would be affected by training data and hyperparameters. In addition, the intra-individual variability of the macro mode cannot be ignored. However, we explored the robustness of macro mode and gave a feasible selection rule for macro mode variability problem.

In PathFinder, the macro mode can achieve state-of-the-art prognostic performance as micro mode does. Considering that numerous studies have achieved multi-class tissue segmentation across various cancer types^38,39^, further exploration of the impact of these ready-made segmentation maps on prognosis may lead to new discoveries. Moreover, benefiting from its simple and easy-to-use features, PathFinder can be easily migrated to similar tasks such as spatial multi-omics and three-dimensional pathological prognosis to discover new biomarkers in different modalities^40–42^. We expect Pathfinder as a fundamental mechanism to better integrate the two fields of clinical prognosis and AI, and inspire more meaningful discoveries.

## Methods

### Evaluation Details and Statistical Analysis

The multi-class classification performance of PaSegNet was evaluated on the internal test set of QHCG and external test sets of TCGA and PAIP. Macro-average accuracy of multi-class classification was calculated based on the confusion matrices of test sets. Nonparametric bootstrapping with 1,000 samples was used to compute 95% confidence intervals of macro-average AUC.

The predicted risk scores for MacroNet, MicroNet and M2MNet were first evaluated on the same validation splits of TCGA in a 10-fold cross validation. Cross-validated C-Index performance is reported as the media C-Index over the 10-folds. The significance level was determined using a two-sided Mann-Whitney-Wilcoxon test. The prognostic performances of MacroNet, MicroNet and M2MNet trained on TCGA were then evaluated on QHCG dataset, and nonparametric bootstrapping with 1,000 samples was used to compute 95% confidence intervals of C-Indices. The significance level was determined using a two-sided two-sample *t*-test. For significance testing of patient stratification in Kaplan-Meier analysis, we used the logrank test to measure if the difference of two survival distributions was statistically significant (P-Value < 0.05), and the cutoff value of Kaplan-Meier analysis was selected as the value with smallest P-Value. Univariable and multivariable analyses of factors associated with prognosis were selected to test whether the risk scores were independent prognostic indicators.

The quantified new biomarkers of NEC and TND were first evaluated on TCGA and QHCG datasets by using Kaplan-Meier analysis and logrank test to measure whether the biomarkers can stratify two survival distributions significantly. Univariable and multivariable analyses were further used to test whether the new biomarkers were independent prognostic indicators.

### Computational Hardware and Software

Python (version 3.7.9) packages used by the project include PyTorch (version 1.8.0), Lifelines (version 0.25.11), NumPy (version 1.19.2), Pandas (version 1.2.2), Albumentations (version 0.5.2), OpenCV (version 4.5.1), Pillow (version 7.2.0) and OpenSlide (version 1.1.2). All WSIs were processed on Intel Xeon multi-core CPUs (Central Processing Units) and a total of four 3090 GPUs (Graphics Processing Units). Deep learning models were trained with Nvidia softwares CUDA 11.1 and cuDNN 8.0.5. Saliency was implemented using Captum (version 0.2.0). Statistical analyses such as two-sampled *t*-tests used implementations from SciPy (version 1.4.1), and logrank tests, univariable and multivariable analyses used implementations from Lifelines (version 0.25.11). Plotting and visualization packages were generated using Seaborn (version 0.9.0) and Matplotlib (version 3.1.1).

## Data availability

The data that supports the plots within this paper and other findings of this study will be available after the article is published.

## Code availability

The codes that support the findings of this study will be available after the article is published.

## Acknowledgements

We thank Y. Gao, S. Yang and X. Chen for helpful comments on the manuscript.

## Authors contributions

L.K. and J.L. conceived the idea. L.K. and H.Y. supervised the project. J.L. and Y.X. performed the experiments. Y.X., Y.J., and W.M. curated the QHCG dataset. J.L., Y.X., and W.Z. analyzed the results.

Q.D. provided helpful discussions on the project design. J.L. and L.K. prepared the manuscript with inputs from all co-authors.

